# HERC2 promotes BLM and WRN to suppress G-quadruplex DNA

**DOI:** 10.1101/341420

**Authors:** Wenwen Wu, Nana Rokutanda, Jun Takeuchi, Yongqiang Lai, Reo Maruyama, Yukiko Togashi, Hiroyuki Nishikawa, Naoko Arai, Yasuo Miyoshi, Nao Suzuki, Yasushi Saeki, Keiji Tanaka, Tomohiko Ohta

## Abstract

BLM and WRN are RecQ DNA helicases essential for genomic stability. Here we demonstrate that HERC2, a HECT E3 ligase, is critical for their functions to suppress G-quadruplex (G4) DNA. HERC2 interacts with BLM, WRN, and replication protein A (RPA) complexes during S-phase of the cell cycle. Depletion of HERC2 dissociates RPA from BLM and WRN complexes and significantly increases G4 formation. Triple depletion revealed that HERC2 has an epistatic relationship with BLM and WRN in their G4- suppressing function. *In vitro*, HERC2 releases RPA onto single-stranded DNA (ssDNA), rather than anchoring onto RPA-coated ssDNA. CRISPR/Cas9-mediated deletion of the catalytic ubiquitin-binding site of HERC2 causes RPA accumulation in the helicase complexes and increases G4, indicating an essential role for E3 activity in G4 suppression. Both HERC2 depletion and E3 inactivation sensitize cells to the G4-interacting compounds, telomestatin and pyridostatin. Overall, HERC2 is a master regulator of G4 suppression and affects the sensitivity of cells to G4 stabilizers.

## INTRODUCTION

Specific guanine-rich DNA motifs containing four stretches of three or more consecutive guanines can fold into a secondary structure, known as the G-quadruplex (G4), via Hoogsteen base pairing stabilized by a monovalent cation [1, 2]. G4 structures were originally characterized *in vitro* using biophysical techniques, and recent high-throughput sequencing techniques have identified more than 700,000 G4 structures genome-wide [3]. Accumulating evidence has demonstrated its important functional roles *in vivo* that have recently attracted attention. G4 plays critical roles in telomere maintenance, replication initiation, and transcriptional initiation and termination. On the other hand, G4 can constitute obstacles for replication fork progression and transcription. Fork stalling and transcription-associated DNA damage caused by G4s raise the potential to generate genomic instability and consequent development of cancer or disease [1, 2, 4]. Clinically, G4s may represent molecular targets to exploit genomic instability in cancers using G4- interacting compounds, such as telomestatin and pyridostatin [4-6].

Several ATP-dependent DNA helicases have been shown to disassemble G4 [7]. Among these, RecQ helicase family members, including BLM and WRN, that unwind noncanonical DNA structures in a 3′ to 5′ direction [8], play major roles in G4 suppression [7, 9, 10]. Deficiencies in BLM and WRN cause Bloom and Werner syndromes, respectively; rare autosomal recessive disorders categorized in chromosome instability syndrome. Bloom syndrome is characterized by growth retardation, immunodeficiency, hypersensitivity to sunlight, and predisposition to a wide spectrum of cancers [11]. Cells from Bloom syndrome patients show defects in DNA replication and homologous recombination, and exhibit a high frequency of sister chromatid exchanges [11]. In addition to G4s, the DNA substrates unwound by BLM include 3′-tailed duplexes, bubble structures, forked duplexes, DNA displacement loops, and double Holliday Junctions (dHJs) [8, 12, 13]. BLM is critical for repairing replication fork restoration by unwinding noncanonical DNA structures, preventing inappropriate recombination, and resetting stalled forks by fork regression [14, 15]. Werner syndrome is an adult-onset syndrome with a range of features consistent with accelerated aging, including short stature, diabetes mellitus, osteoporosis, atherosclerosis, malignancies, and early death. [16]. Similar to BLM, WRN protein plays critical roles in DNA repair, replication, transcription, and telomere maintenance, and disrupts G4s, bubble structures, and dHJs at perturbed replication forks or DNA double-strand breaks [7, 8, 17–19].

Both BLM and WRN physically interact with replication protein A (RPA) ([20, 21], a heterotrimeric complex of RPA1 (RPA70), RPA2 (RPA32), and RPA3 (RPA14) that binds to single-stranded DNA (ssDNA) and prevents its spontaneous annealing. The interaction significantly stimulates helicases activity to unwind long duplex DNAs [20–22], and accumulating evidence supports a function of RPA in G4 unfolding [23, 24]. In addition to helicase activity, both BLM and WRN possess strand annealing activity, and rewind ssDNAs after unwinding double-stranded DNA (dsDNA) [25]. RPA increases the helicase activity, but inhibits their strand annealing activity [20–22]. Similarly, BLM-mediated G4 unfolding is followed by G4 refolding [26]. Therefore, the helicases may require RPA to fulfill its function to suppress G4 *in vivo*. However, regulation of RPA–helicase interaction *in vivo* is not well understood.

HERC2 is a large HECT and RCC-like domain-containing protein comprising 4834 amino acids. Mutations in the *HERC2* gene have been linked to a developmental delay [27, 28], and mice with homozygous truncation of the gene are not viable [29]. While HERC2 performs a range of functions with various interacting proteins, one of its major roles is the regulation of DNA replication and damage response. HERC2 controls nucleotide excision repair [30] and homologous recombination [31], fine-tunes ubiquitin-mediated DNA damage response via the deubiquitinating enzyme, USP16 [32], and modulates p53 activity [33]. HERC2 interacts with BRCA1 and is capable of suppressing G_2_–M checkpoint activity by degrading BRCA1 in BARD1-depleted cells [34]. HERC2 is a component of the replication fork complex [35] and suppresses the Chk1-directed checkpoint by degrading the deubiquitinating enzyme, USP20 [36, 37]. Hence, while HERC2 likely plays a pivotal role in DNA replication and the damage response, its function remains unclear. Here, we demonstrate a previously unrecognized function of HERC2 that involves novel binding partners BLM, WRN, and RPA. HERC2 suppresses G4 via BLM and WRN, in a process reliant in the ubiquitin ligase activity of HERC2. Inhibition of HERC2 function sensitizes cells to G4-interacting compounds. Interestingly, HERC2 expression is frequently reduced in many types of cancers; therefore, these results highlight the critical role of HERC2 on chromosomal stability in cancers, as well as its importance as an indicator for the efficiency of G4 stabilizers in cancer treatments.

## RESULTS

### HERC2 interacts with BLM, WRN, and RPA complexes in S-phase nuclear extracts

To investigate the function of HERC2, we raised rabbit polyclonal antibody specific to a C-terminal fragment of HERC2 and analyzed the HERC2 immunocomplex isolated from HeLa and HCT116 cells using mass spectrometry. In addition to the proteins previously reported to interact with HERC2 (Figure S1A), mass spectrometry analyses identified all subunits of the BTR complex consisting of <u>B</u>LM, topoisomerase IIIa (<u>T</u>OP3A), <u>R</u>MI1 (also known as BLAP75), and <u>R</u>MI2 (BLAP18), that function as a dissolvasome to dissolve dHJs and other secondary DNA structures [38–40] (Figure 1A). The analyses also identified WRN, all subunits of the RPA complex, and another BLM-interacting protein, BRCA1/BARD1 [41]. HERC2 immunoprecipitation followed by immunoblotting verified these interactions (Figure 1B).

**Figure 1.**
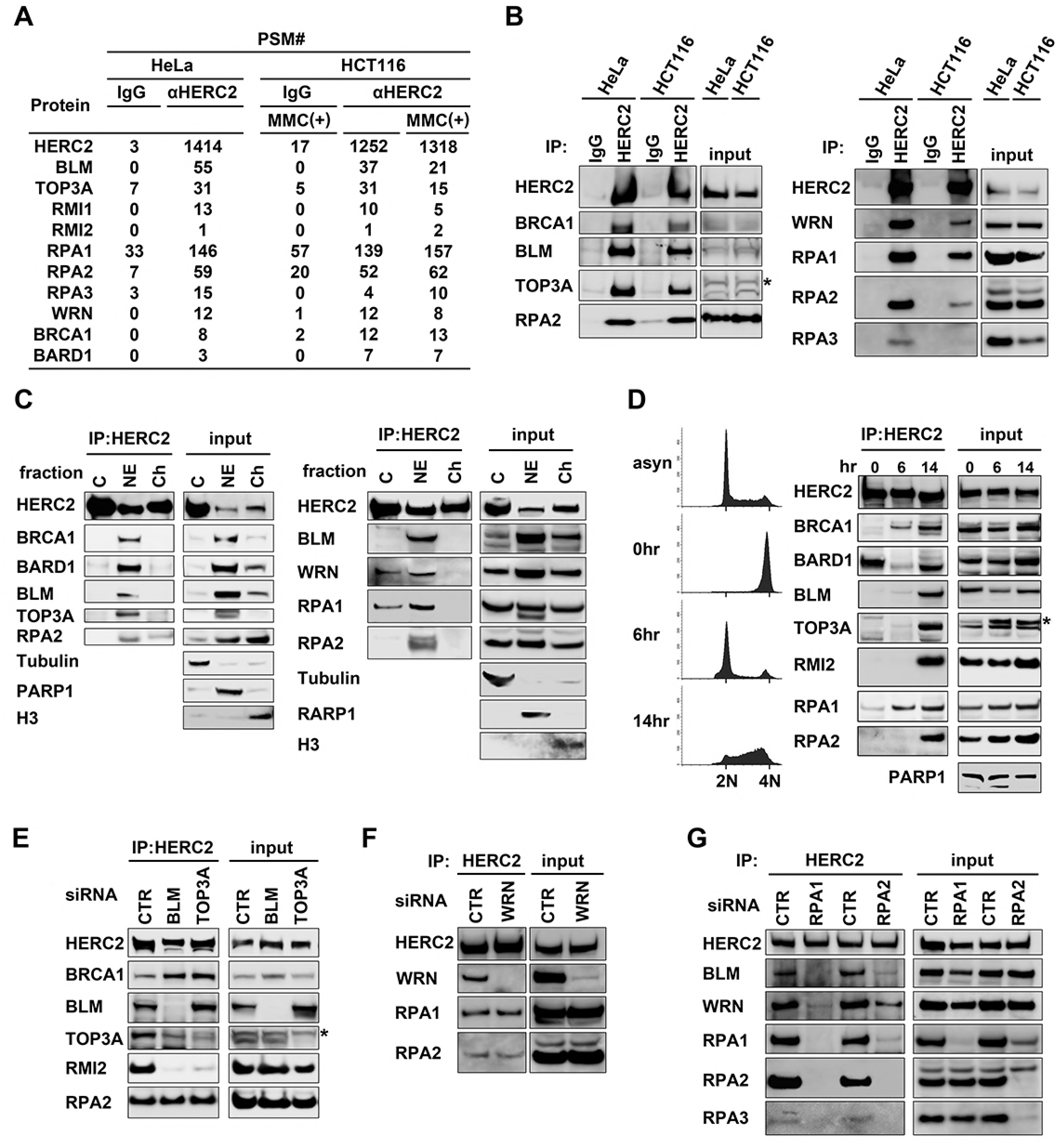
HERC2 interacts with BLM, WRN, and RPA complexes in S-phase nuclear fractions. **(A)** Affinity purification of immunocomplexes from HeLa or HCT116 cells treated with or without mitomycin C (MMC) was performed using an antibody to HERC2 or preimmune IgG coupled with magnetic beads. Lists of the BLM-related proteins identified via mass spectrometry analysis are shown (PSM, peptide spectrum matches). **(B)** Lysates from HeLa and HCT116 cells were immunoprecipitated (IP) with anti-HERC2 antibody or IgG followed by immunoblotting with the indicated antibodies. Inputs were also loaded. **(C)** Cytosolic (C), soluble nuclear extract, and solubilized chromatin (Ch) fractions were prepared from HeLa cells and immunoprecipitated with anti-HERC2 antibody followed by immunoblotting. **(D)** HeLa cells were synchronized with thymidine-nocodazole block, released, and harvested at the indicated time points. Synchronization was monitored by flow cytometry (left panels). Nuclear extracts were subjected to immunoprecipitation with anti-HERC2 antibody followed by immunoblotting. asyn, asynchronous cells. **(E**–**G)** HeLa cells were transfected with either control siRNA (CTR) or siRNAs specific to BLM, TOP3A (E), WRN (F), and RPA1/2 (G) and subjected to immunoprecipitation with anti-HERC2 antibody followed by immunoblotting. Asterisk denotes a nonspecific band.

HERC2 is a nuclear–cytoplasmic shuttle protein that localizes in both fractions and is abundantly expressed during all phases of the cell cycle [30, 34]. To uncover specific cellular conditions in which HERC2 functions with BLM, WRN, and RPA, we analyzed their interaction in separated cellular fractions and in each phase of the cell cycle. Fractionation analysis revealed that HERC2 interacts with these complexes mostly in soluble nuclear extracts but not in cytosolic and chromatin fractions (Figure 1C). Fractionation efficiency was demonstrated with tubulin (cytosol), PARP1 (nuclear extract), and histone H3 (chromatin). We next synchronized HeLa cells. To minimize the effects according to cellular response to thymidine-induced replication stress, cells were treated with a thymidine-nocodazole block and released into the cell cycle, then harvested at mitosis (0 h), G_1_ (6 h), and S (14 h) phases. HERC2 immunoprecipitation from nuclear extracts followed by immunoblotting demonstrated that HERC2 interacts with the BTR, WRN, RPA, and BRCA1/BARD1 complexes mainly during S-phase (Figures 1D and S1B).

Both BLM and WRN are capable of directly interacting with RPA *in vitro* [20–22], suggesting that the HERC2–RPA interaction is mediated by BLM and/or WRN, or vice versa. To clarify the configuration of the HERC2 complex, we inhibited subunits of the complex using siRNA and performed HERC2 immunoprecipitation followed by immunoblotting. Depletion of BLM clearly suppressed coprecipitation of TOP3A and RMI2, whereas depletion of TOP3A only inhibited coprecipitation of RMI2 but not BLM (Figure 1E), indicating that BLM bridges the interaction between the HERC2 and BTR complex. In contrast, interaction of HERC2 with RPA1 and RPA2 was not inhibited by the depletion of either BLM or TOP3A (Figures 1E and S1C). The HERC2–RPA interaction was not inhibited by depletion of WRN (Figure 1F), indicating that RPA interacts with HERC2 independent of BLM and WRN. We next investigated depletion of RPA1 and RPA2. Depletion of RPA2 significantly reduced the steady state level of RPA1 and RPA3, whereas depletion of RPA1 did not dramatically affect that of RPA2 and RPA3 (Figure 1G), as previously reported [42]. In the condition depletion of RPA1 inhibited HERC2–RPA2 and HERC2–RPA3 interactions (Figure 1G), suggesting that RPA1 mediates the interaction of HERC2 with the RPA complex. In addition, both RPA1 and RPA2 depletion significantly reduced the level of BLM and WRN coprecipitated with HERC2, suggesting that RPA1 is critical for HERC2–BLM–RPA or HERC2–WRN–RPA complex formation. Although BLM interacts with BRCA1 [41], depletion of BLM or BRCA1 did not inhibit HERC2–BRCA1 or HERC2–BLM interactions, respectively (Figures 1E and S1D), suggesting that BRCA1 and BLM independently interact with HERC2. Although BLM forms a BRAFT complex with Fanconi anemia (FA) core complex [43], FA proteins were not detected by mass spectrometry analyses of HERC2 immunocomplexes, suggesting that the complex interacting with HERC2 is not the BRAFT complex.

### HERC2 mediates BLM–RPA and WRN–RPA interactions in unstressed cells

The interaction between HERC2 and BLM/WRN/RPA prompted us to investigate whether HERC2 affects BLM–RPA and WRN–RPA complex formation. To test this, we established HeLa cells harboring doxycyclin (Dox)-inducible shRNA against HERC2 (hereafter referred to as HeLa-shHERC2 cells). Dox addition induced efficient depletion of HERC2 in the cells (Figure 2A). Remarkably, anti-BLM immunoprecipitation followed by anti-RPA immunoblotting revealed that HERC2 depletion significantly reduced the interaction between BLM and RPA (Figure 2A). The same result was also observed in HCT116 cells with HERC2 knockdown (Figure S2A).

**Figure 2.**
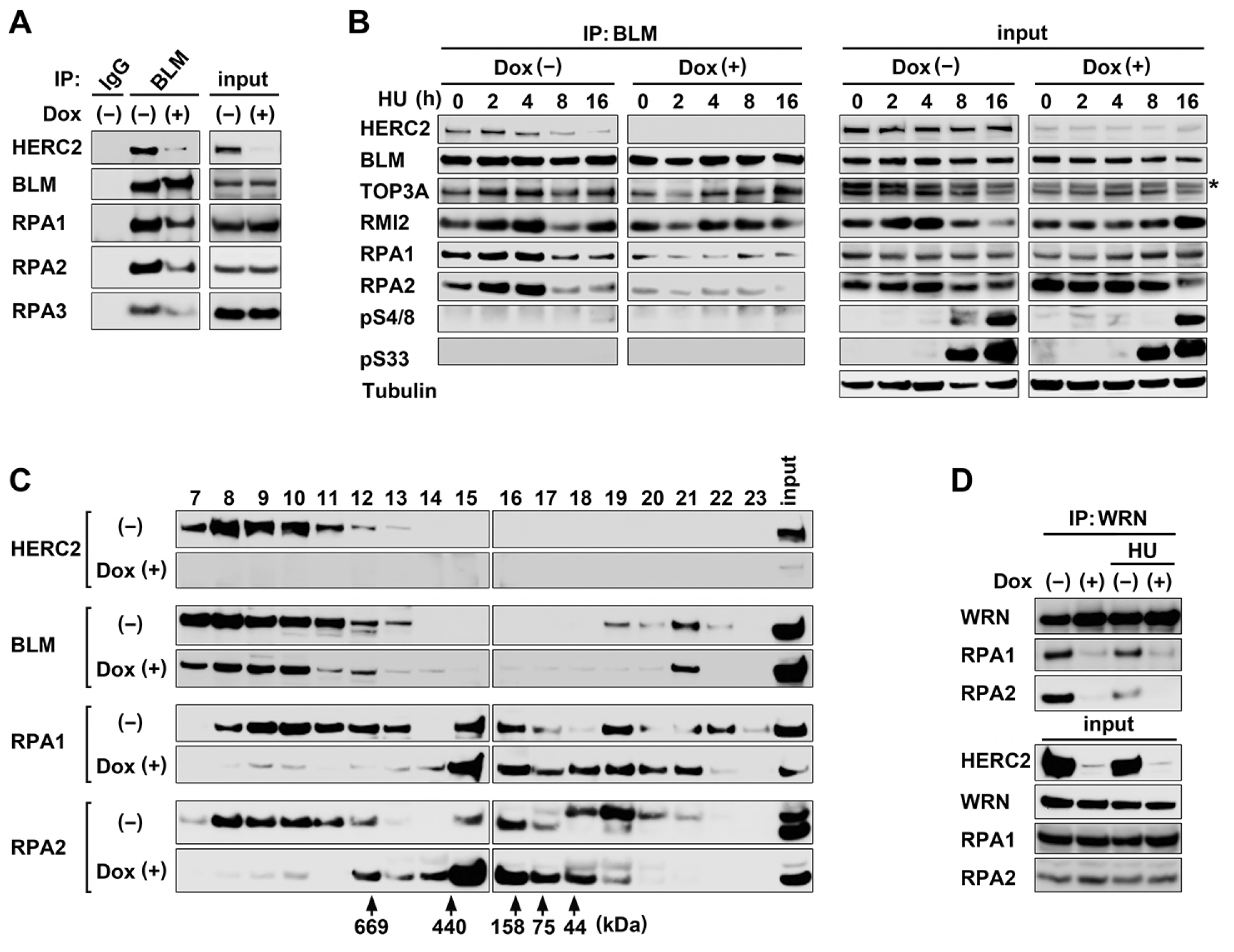
HERC2 regulates BLM–RPA and WRN–RPA complex formation. **(A)** HeLa-shHERC2 cells were induced or not with Dox and lysates were immunoprecipitated (IP) with anti-BLM antibody or IgG followed by immunoblotting with the indicated antibodies. Inputs were also loaded. **(B)** HeLa-shHERC2 cells were induced or not with Dox and treated with hydroxyurea (HU, 1 mM) for the indicated time, and were subjected to anti-BLM immunoprecipitation followed by immunoblotting. Asterisk denotes a nonspecific band. **(C)** HERC2 depletion alters the molecular weight of the RPA complex. HeLa-shHERC2 cells were induced or not with Dox. Nuclear extracts were fractionated by size-exclusion chromatography on a Superose 6 gel filtration column, and the indicated proteins were detected by immunoblotting. The elution positions of the protein standards are indicated at the bottom. **(D)** HeLa-shHERC2 cells were induced or not with Dox and lysates were subjected to anti-WRN immunoprecipitation followed by immunoblotting with the indicated antibodies. HU (1 mM) was added 16 h prior to harvesting, where indicated.

During the course of the experiments, we noticed that the HERC2-dependent BLM–RPA interaction was reduced by replication stress generated by 16 h exposure to mitomycin C (MMC) (Figure S2B). To further analyze the effects of replication stress on BLM–RPA complex formation in the presence or absence of HERC2, we analyzed a time course after the stress. HeLa-shHERC2 cells were induced or not with Dox and incubated with either hydroxyurea (HU) or MMC then subjected to anti-BLM immunoprecipitation followed by immunoblotting (Figure 2B and S2C). Without HERC2 depletion, the interaction between BLM and RPA1/RPA2 was slightly enhanced at 2 h after the addition of HU or MMC then significantly reduced at 8 and 16 h. The interaction between BLM and HERC2 also reduced after the stress. On the contrary, in HERC2-depleted cells the BLM–RPA interaction remained significantly reduced during the time course. Phosphorylation of RPA2 at Ser 4/8 and Ser33, which are induced by DNA-dependent protein kinase and ATR [44], was increased at 8 or 16 h after the stress when the BLM– RPA interaction was reduced, indicating that RPA2 interacted with BLM and affected by HERC2 is in its unphosphorylated form. Together, these findings indicate that HERC2 mediates BLM–RPA interaction in unstressed cells.

To further confirm the effects of HERC2 on BLM–RPA complex formation, we performed gel filtration analyses using HeLa-shHERC2 cells (Figure 2C). Without HERC2-depletion, RPA1 and RPA2 coeluted with BLM in high-molecular-mass fractions at fraction 8 to 13 where HERC2 also coeluted, consistent with the complex formation of these proteins. Notably, the RPA1 and RPA2 fractions shifted dramatically to lower molecular weights with elution peaks around 158 to 440 kDa (fractions 15 and 16) following Dox induction, whereas BLM remained in the high-molecular-mass fractions, supporting BLM–RPA dissociation. These results indicate that RPA exists in highmolecular-mass complex(es) with BLM in a HERC2-dependent manner. Gel filtration experiments showed that distributions of fractions for BLM and RPAs following MMC treatment were analogous to that induced by HERC2 depletion (Figure S2D), supporting a role for HERC2 in BLM–RPA interaction in unstressed cells.

The HERC2-dependent interaction of BLM and RPA prompted us to test whether HERC2 was also involved in WRN–RPA interaction in the same way. As shown in Figure 2D, HERC2 was found to mediate WRN–RPA interaction in unstressed cells, and this interaction was reduced after a 16-h incubation with HU. Together these results suggest that HERC2 does not simply interact with BLM–RPA and WRN–RPA complexes, but is also required for their formation.

### HERC2 is epistatic to BLM and WRN in G4 suppressing function

RPA plays a supportive role in the helicase reaction of BLM and WRN *in vitro* [20–22]; therefore, the absence of RPA in the BLM and WRN complexes by HERC2 depletion may affect the biological function of the helicases. To clarify this, we investigated whether HERC2 depletion resulted in G4 accumulation, a common phenotype shared by BLM and WRN dysfunction [7, 9, 10]. G4 was analyzed by immunostaining using a G4-specific antibody BG4 [45]. The antibody detected G4 as nuclear foci that were enhanced by a BLM inhibitor or G4 stabilizers (Figures S3A and S3B). HeLa-shHERC2 cells were then induced or not with Dox and immunostained for G4 (Figure 3A). Without replication stress, G4 accumulation was remarkably enhanced by HERC2 depletion, indicating that HERC2 represses constitutive G4 accumulation. Brief exposure to HU reduced G4, consistent with previous observations that replication arrest via brief exposure to aphidicolin reduced G4 [45]. HERC2-depleted cells briefly exposed to HU still exhibited significantly higher rates of G4, reflecting an accumulation at the basal level. Similar results were also observed in HERC2-depleted U2OS cells (Figure S3C).

The similarities in phenotype shared by HERC2 depletion with dysfunction of BLM and WRN prompted us to examine whether the observed effects of HERC2 depletion on G4 were mediated by BLM and/or WRN dysfunction. To examine this, we analyzed the epistatic effects of HERC2 and BLM/WRN depletion. HeLa-shHERC2 cells were transfected with control siRNA or BLM-specific siRNA then induced or not with Dox and immunostained for G4 (Figure 3B). The efficient depletion of both proteins was confirmed by immunoblotting. To our surprise, the level of G4 accumulation in HERC2-depleted cells was much higher than in BLM-depleted cells. In addition, BLM depletion in HERC2-depleted cells did not change the level of G4 accumulation compared with that of cells with HERC2-single depletion, suggesting that HERC2 is epistatic to BLM in the suppression of G4. An epistatic relationship was also observed between HERC2 and WRN (Figure 3C).

Finally, we performed triple knockdown of HERC2, BLM, and WRN and analyzed their epistatic relationship during G4 regulation (Figure 3D). The efficient depletion of all three proteins was confirmed by immunoblotting. G4 accumulation in cells with a single BLM or WRN depletion was similar. On the other hand, double depletions of BLM and WRN caused significantly higher levels of G4 than a single depletion, suggesting that BLM and WRN function in different pathways or influence different subsets of G4. Interestingly, G4 accumulation in BLM/WRN double-depleted cells was approximately equal to that of HERC2-depleted cells, and a BLM/WRN double depletion did not further increase G4 in HERC2-depleted cells. Together, these results indicate that HERC2 suppresses G4 in a manner epistatic to the additive effects of BLM and WRN, the two major RecQ helicases involved in G4 unwinding.

**Figure 3.**
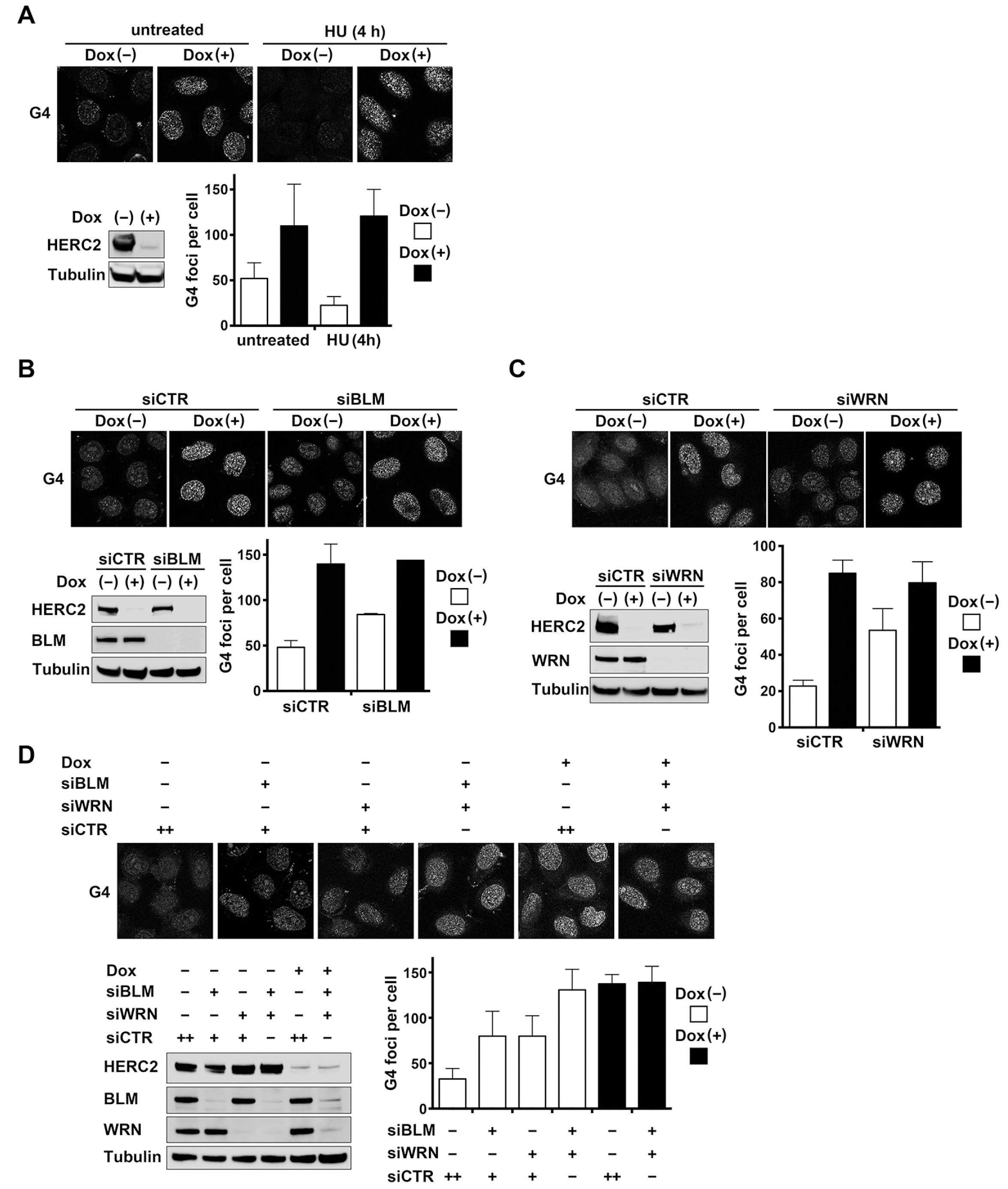
HERC2 depletion increases G4 formation. **(A)** HeLa-shHERC2 cells were induced or not with Dox, and subjected to immunoblotting with the indicated antibodies (left lower panel), or treated or not with HU (200 μM) for 4 h and immunostained for G4 (upper panel). Quantification of the mechanically counted G4 foci per cell are shown (right lower panel). Error bars represent standard error of the mean (SEM) of duplicate experiments, each based on more than 100 cells. **(B**–**D)** HeLa-shHERC2 cells were transfected with either control siRNA (CTR) or siRNAs specific to BLM (B), WRN (C), or BLM, and WRN (D) as indicated, induced or not with Dox, and subjected to immunoblotting with the indicated antibodies (left lower panels) or immunostained for G4 (upper panels). Quantification of the mechanically counted G4 foci per cell is shown (right lower panels). Error bars represent SEM of duplicate experiments, each based on more than 100 cells.

### HERC2 does not anchor to RPA-coated ssDNA

In addition to its role to prevent the secondary structure of DNA, ssDNA-binding RPA plays a critical role as a scaffold for DNA metabolism proteins such as ATR–ATRIP in the stalled replication fork in response to replication stress or DNA damage [46]. Therefore, one obvious possibility for the role of HERC2, which mediates interaction of BLM, WRN, and RPA, is recruitment of the helicases to the RPA scaffold. However, this was contradicted by the observation that HERC2 facilitated the interaction in unstressed conditions that reduced with ATR-mediated phosphorylation of RPA in response to replication stresses (Figures 2B and S2C). To clarify this, we first examined the intracellular localization of HERC2. HERC2 localizes both in the nucleus and the cytoplasm [30, 34] and colocalizes with replication fork complex proteins including PCNA in the nucleus [35]. Costaining of HERC2 and RPA2 showed that HERC2 colocalizes with RPA2 in dispersed nuclear foci in normal proliferating cells (Figure 4A, upper panels). This colocalization disappeared as RPA2 became discrete nuclear foci after a 16-h incubation with HU (lower panels), suggesting that HERC2 does not interact with RPA-coated ssDNA at stalled replication forks. The specificity of the HERC2 antibody used for immunostaining was confirmed by depletion of HERC2 (Figure S4).

**Figure 4.**
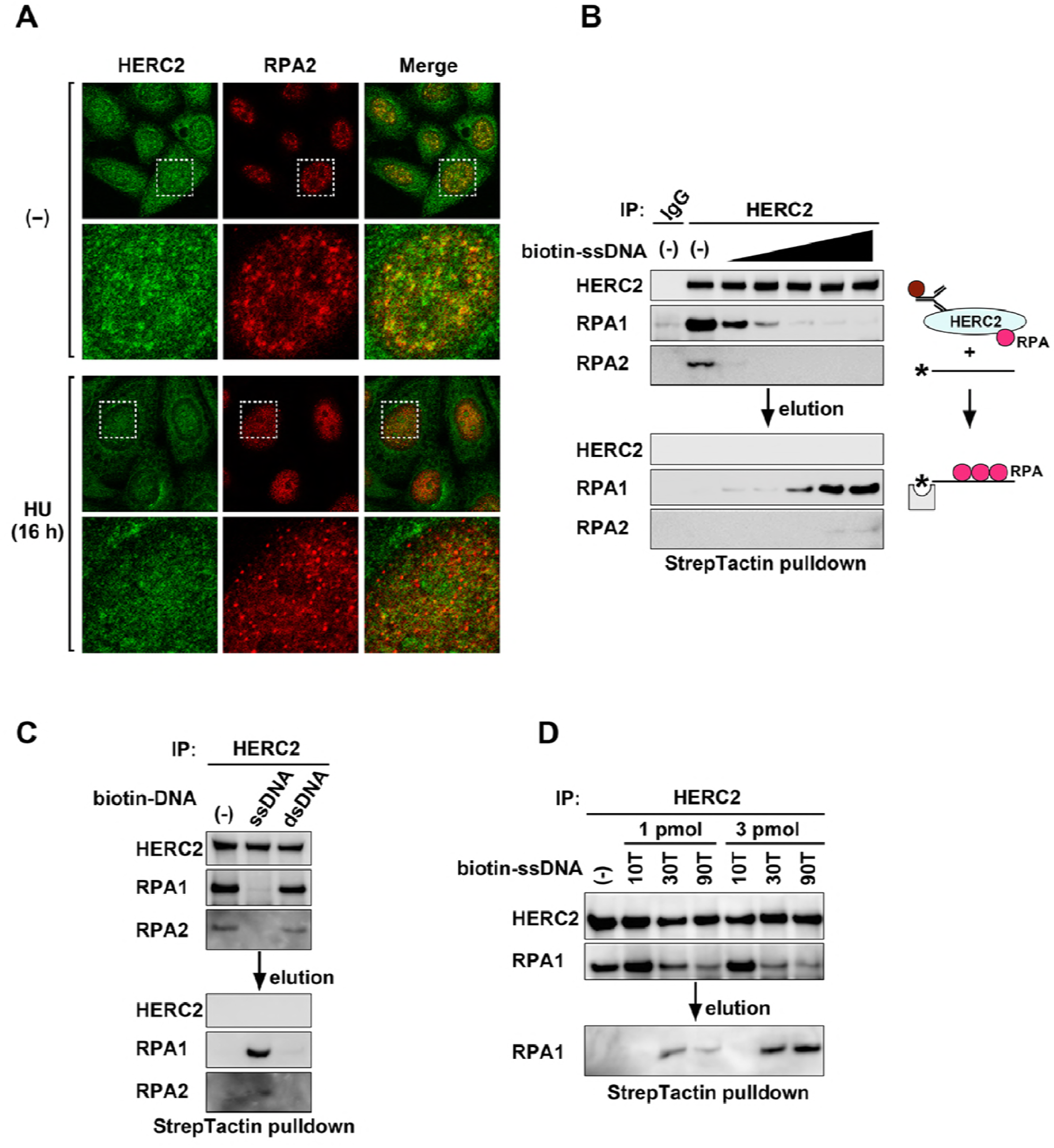
HERC2 does not anchor to RPA-coated ssDNA. **(A)** HERC2 colocalizes with RPA2 in the nuclear without replication stress. HeLa cells were treated or not (-) with HU (1 mM) for 16 h, and immunostained with the indicated antibodies. Each lower panel shows one nucleus (dashed line) that has been magnified. **(B)** HERC2 complex from HeLa cell lysate immunoprecipitated with control IgG or anti-HERC2 antibody-coated dynabeads were incubated with an increasing amount (0, 0.1, 0.3, 1, 3, 10 pmol) of biotin-labeled ssDNA. Elution was subjected to StrepTactin-pulldown. The StrepTactin-bound proteins (lower panels) and proteins bound to the remained dynabeads (upper panels) were immunoblotted with the indicated antibodies. A schematic representation of the method is shown. Asterisk indicates biotin, bar indicates ssDNA, and gray square indicates StrepTactin. **(C and D)** HERC2 immunocomplex was subjected to biotin-labeled DNAs as in (A) except that 10 pmol of ssDNA and dsDNA (C) or ssDNA with different length and different amount (D) were used.

To further clarify the configuration of the HERC2–RPA–ssDNA interaction, we examined whether the HERC2–RPA complex was capable of interacting with ssDNA *in vitro*. HERC2 complexes were immunoprecipitated from HeLa cell lysates using anti-HERC2 antibody-coated dynabeads, and incubated with biotin-labeled ssDNA (Figure 4B). Notably, addition of ssDNA removed RPA from HERC2 in dose-dependent manner. We then subjected the elution to StrepTactin-pulldown and confirmed that RPA was transferred from HERC2 to ssDNA. The transfer was ssDNA specific, but not specific for dsDNA (Figure 4C), and also required a certain length (∼ 30 nt) of ssDNA (Figure 4D). Thus, the results indicate that HERC2 is not scaffold to mediate protein interactions on RPA-coated ssDNA. Instead, HERC2 may facilitate the delivery of RPA to ssDNA.

### Deletion of the catalytic site of HERC2 causes RPA accumulation in BLM and WRN complexes and increases G4

HERC2 comprises Cys4762, a conserved catalytic ubiquitin-binding site of the HECT domain in most C-termini that is essential for the enzyme activity [34]. To uncover whether the E3 ubiquitin ligase activity affects BLM–RPA complex formation and is involved in the observed phenotypes of HERC2 dysfunction, we disrupted the C-terminal end of the HERC2 gene using CRISPR/Cas9 nuclease-mediated genome editing and established a HCT116 cell line with a homozygous insertion mutation that generates a premature stop codon at Glu4758 and deletes Cys4762 (hereafter referred to as HERC2^ΔE3/ΔE3^, Figure 5A). The steady state level of mutated HERC2 was the same level as the wild type (WT), as determined by an antibody to the epitope at amino acids 1781–1974 of HERC2 (Figure 5B). On the contrary, an antibody to the epitope at amino acids 4784–4834 failed to detect the mutant HERC2, indicating successful deletion of the HERC2 C-terminus.

**Figure 5.**
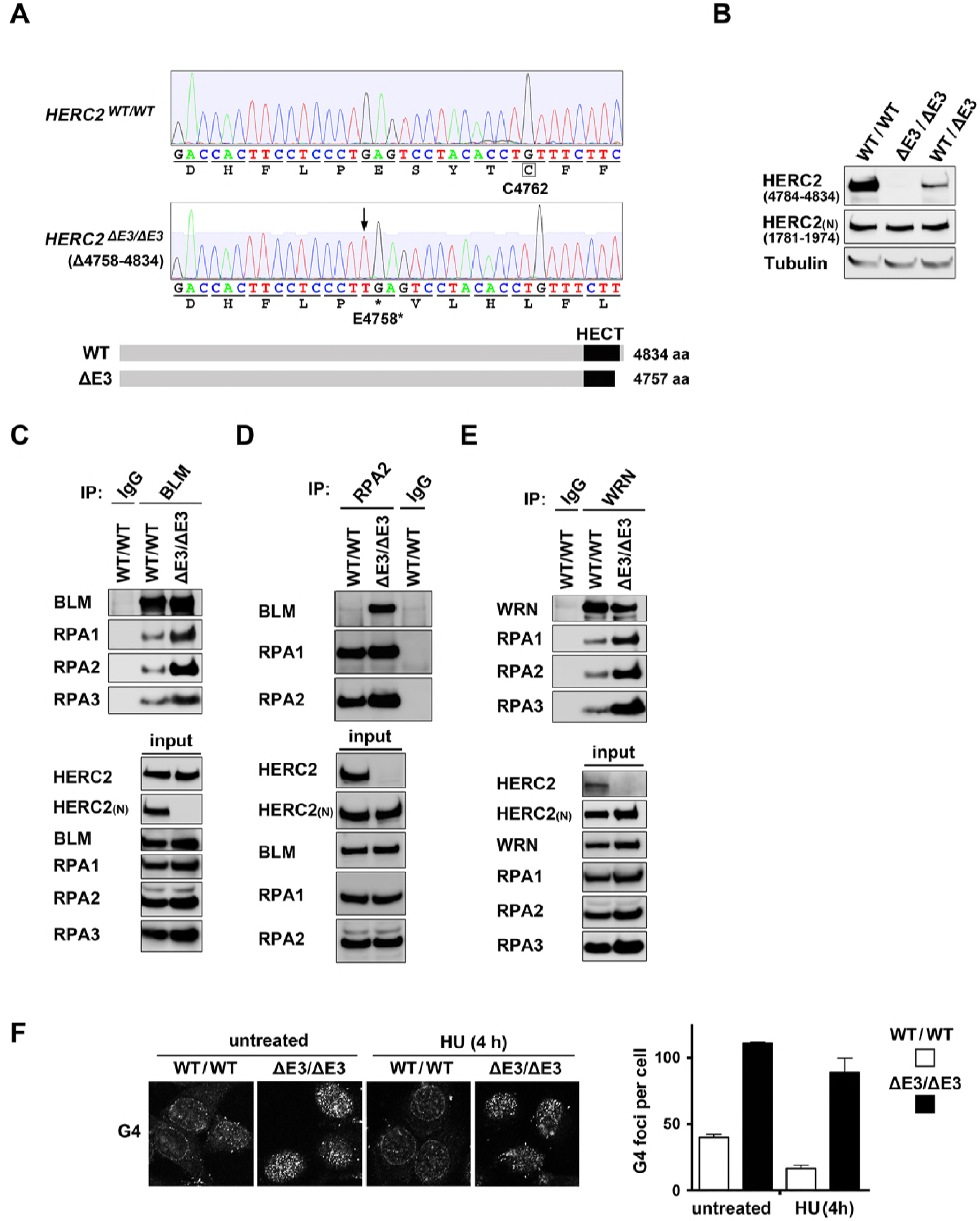
CRISPR/Cas9-mediated deletion of the ubiquitin-binding site of HERC2 increases G4. **(A)** CRISPR/Cas9-mediated deletion of the ubiquitin-binding site of HERC2 in HCT116 cells. Sequencing chromatogram of genomic DNA shows the region with an insertion (arrow) in targeted HCT116 cell line (HERC2^ΔE3/ΔE3^). The mutation causes premature stop codon at E4758 that eliminates the catalytic site C4762. Schematic representation of WT and Δ E3 HERC2 proteins are shown below. **(B)** Lysates from HCT116 cells with WT, homozygous (HERC2^ΔE3/ΔE3^), and heterozygous (HERC2^WT/ΔE3^) mutation were subjected to immunoblotting with antibody to epitope 4784–4834 or 1781–1974 (N) of HERC2. **(C to E)** Accumulation of RPA in BLM and WRN complexes via inhibition of HERC2 E3 activity. Cell lysates from WT or HERC2^ΔE3/ΔE3^ cells were subjected to immunoprecipitation with IgG or antibodies to either BLM (C), RPA2 (D), or WRN (E) followed by immunoblotting with the indicated antibodies. Inputs were also loaded. **(F)** Inhibition of HERC2 E3 activity increases G4. WT or HERC2^ΔE3/ΔE3^ cells were treated or not with HU for 4 h and immunostained for G4. Quantification of the mechanically counted G4 foci per cell is shown. Error bars represent SEM of duplicate experiments, each based on more than 100 cells.

Using cells, we first tested whether mutant HERC2 retained the ability to interact with BLM, WRN, and RPA complexes. Due to a lack of an antibody for immunoprecipitation of HERC2^ΔE3/ΔE3^, we performed immunoprecipitation using antibodies to BLM, WRN, RPA1, and RPA2, followed by immunoblotting with a HERC2 antibody to the epitope at amino acids 1781–1974. The results showed that HERC2^ΔE3/ΔE3^ interacts with BLM, WRN, RPA1, and RPA2 at approximately the same level as WT HERC2 (Figure S5A–D). We then tested whether inhibition of HERC2 E3 activity affected BLM–RPA and WRN–RPA complex formations. Interestingly, immunoprecipitation with anti-BLM antibody followed by immunoblotting showed that each subunit of RPA in the BLM immunocomplex was dramatically increased in HERC2^ΔE3/ΔE3^ cells (Figure 5C). Reciprocally, immunoprecipitation with anti-RPA2 antibody flowed by immunoblotting also demonstrated the significant increase of interaction of RPA2 with the BLM in the HERC2^ΔE3/ΔE3^ cells, whereas this antibody barely coimmunoprecipitated with BLM in WT cells (Figure 5D). TOP3A and RMI2, other subunits of BTR, also accumulated in BLM and RPA2 immunocomplexes in unstressed HERC2^ΔE3/ΔE3^ cells (Figure S5E and S5F). Immunoprecipitation with anti-WRN antibody also showed accumulation of the subunits of RPA in the WRN immunocomplex in HERC2^ΔE3/ΔE3^ cells (Figure 5E). Together, these results suggest that E3 activity of HERC2 fine-tunes the amount of the RPA subunits in BTR and WRN complexes. HERC2 may regulate the turnover or stability of the subunit(s) in the complexes via E3 activity.

Since the inhibition of HERC2 E3 activity affected the RPA status in BLM and WRN complexes, we examined whether it impacted the function of the helicases to suppress G4 formation. WT or HERC2^ΔE3/ΔE3^ cells were treated or not with HU for 4 h and immunostained for G4 (Figure 5F). Brief exposure to HU reduced G4 as observed with HeLa cells (Figure 3A). In either condition, G4 accumulation was significantly increased in HERC2^ΔE3/ΔE3^ cells compared with WT cells. These results indicate that HERC2 and its E3 ligase activity are critical for BLM and WRN to suppress G4 accumulation.

### HERC2 dysfunctions sensitize cells to G4-stabilizing compounds

It was reported that depletion of FANCJ, a 5′ to 3′ DNA helicase that also unfolds G4, sensitizes cells to G4-interacting compounds [6]. Thus, accumulation of G4 by HERC2 dysfunction prompted us to test whether it affected the sensitivity of cells to G4-interacting compounds. HeLa-shHERC2 cells were induced or not with Dox, exposed to a range of doses of pyridostatin and telomestatin, the G4-interacting ligands that stabilize G4, and allowed to form colonies. Remarkably, Dox-induced depletion of HERC2 sensitized the cells to both pyridostatin and telomestatin (Figure 6A). Whereas accumulation of G4 may be obstacle for DNA replication, cell growth of the Dox-induced HERC2-depleted cells was only slightly suppressed compared with HERC2-expressing cells (Figure S6A), suggesting that cells were able to overcome the obstacle. In contrast to the G4 stabilizers, HERC2 depletion did not dramatically affect the sensitivity of the cells to replication stress induced by MMC, CPT, and HU (Figure 6A).

We also tested the effect of inhibiting HERC2 E3 activity on the sensitivity of cells to agents using HERC2^ΔE3/ΔE3^ cells. Similar to HERC2-depleted HeLa cells, HCT116 HERC2^ΔE3/ΔE3^ cells exhibited significantly higher sensitivity to the G4-stabilizers than WT cells (Figure 6B). The effect of HERC2 inhibition was especially robust for telomestatininduced cell death. Contrary to HERC-depleted HeLa cells, HCT116 HERC2^ΔE3/ΔE3^ cells grow relatively slowly compared with WT cells although they still can grow continuously (Figure S6B). However, like HERC2-depleted cells, HCT116 HERC2^ΔE3/ΔE3^ cells did not show increased sensitivity to the replication stresses except for that induced by MMC (Figure 6B). Together, these results indicate that HERC2 dysfunction abrogates G4 processing and sensitizes cells to G4 stabilizers.

**Figure 6.**
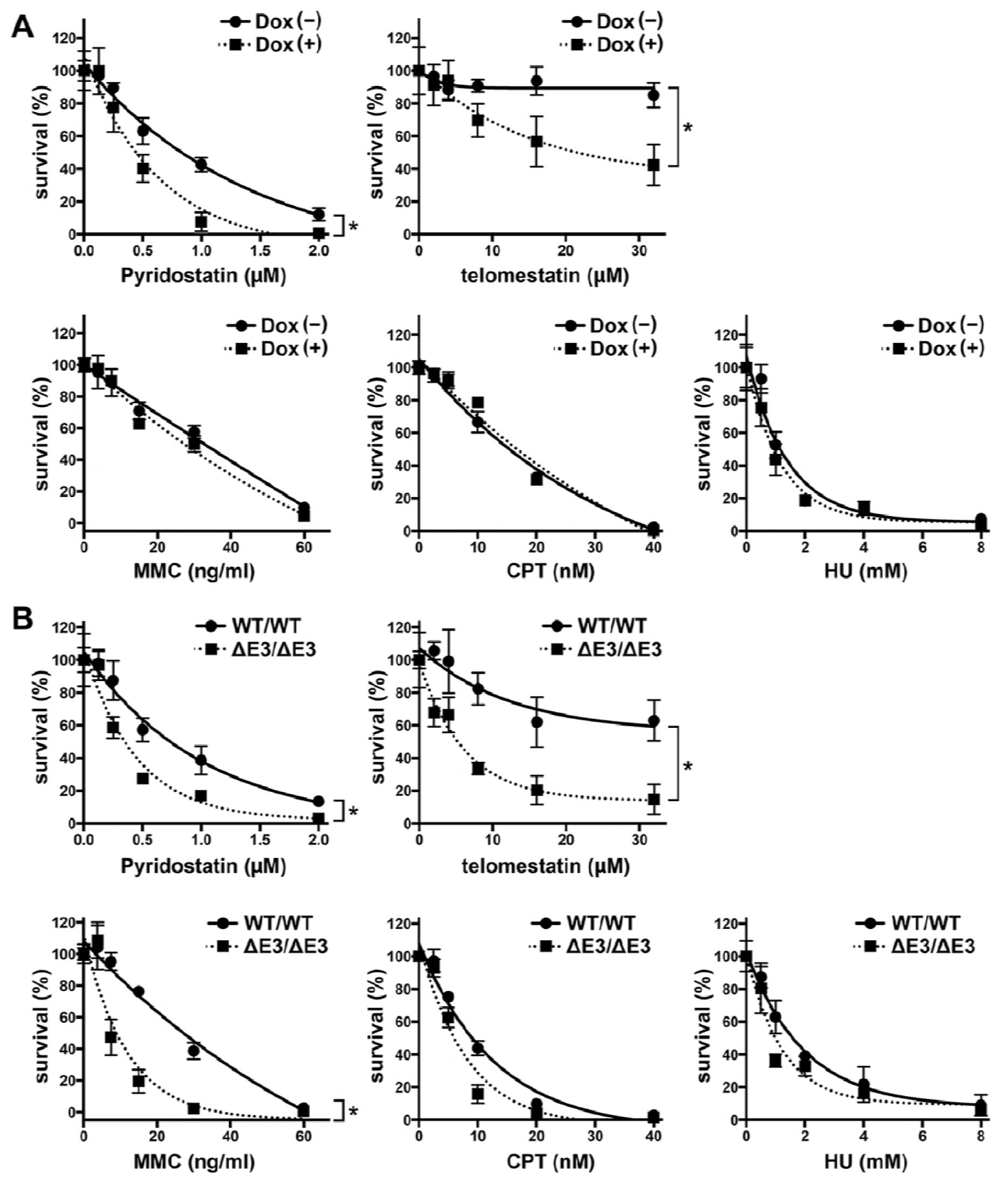
Sensitivity of cells with HERC2 dysfunction to replication stress and G4-stabilizing compounds. **(A and B)** HeLa-shHERC2 cells induced or not with Dox (A), or WT or HERC2^ΔE3/ΔE3^ HCT116 cells (B) were exposed to indicated doses of the chemical agents for 24 h and analyzed for clonogenic survival after two weeks. The data are shown with the nonlinear regression fit curves of one phase decay (GraphPad Prism). Average ± SEM, normalized to cells without agents were derived from triplicate experiments. Statistical significances were calculated using two-way ANOVA. * *P* < 0.000 1.

### HERC2 expression in cancer

To investigate the clinical significance of HERC2 in human cancer, we analyzed TCGA Pan-Cancer clinical data at the UCSC Cancer Genomics Browser. The analysis revealed that the expression of HERC2 is significantly downregulated in various human cancers, including breast invasive carcinoma, bladder urothelial carcinoma, esophageal carcinoma, lung adenocarcinoma, thyroid carcinoma, kidney clear cell carcinoma, and kidney papillary cell carcinoma (Figure S7).

## DISCUSSION

Although initially recognized as a structural interest *in vitro*, G4s have lately emerged as important biological machinery that control many cellular events including gene expression, replication, and telomere lengthening, and affect cell growth, senescence, and genomic instability [1, 2, 4]. Consequently, there is an intense interest in understanding the cellular mechanisms underlying the regulation of G4 formation and removal. In the present study, we provide evidence supporting HERC2 as a master regulator of BLM and WRN, two principal DNA helicases involved in G4 unfolding, to suppress G4 *in vivo* (Figure S8A). HERC2 mediates the interaction of BLM and WRN with RPA, the functional partner of the helicases, in unstressed proliferating S-phase cells. The impact of HERC2 depletion on G4 accumulation is equal to that induced by simultaneous depletion of BLM and WRN, and further depletion of HERC2 does not show an additive effect, indicating an epistatic relationship between HERC2 and these helicases. Inactivation of E3 ligase activity of HERC2 by Cas9/CRISPR-mediated gene editing demonstrated the essential role of the activity to prevent the phenotype. Importantly, dysfunction of HERC2 sensitized cells to G4 stabilizers.

Many previously characterized RPA-interacting proteins were recruited to RPA-coated ssDNA in response to replication stress or DNA damage [46]. Therefore, we hypothesized that HERC2 may act as a scaffold that recruits BLM and WRN to the RPA-coated ssDNA. However, our results demonstrated that this was not the case because (i) HERC2 bridges the helicases and RPA principally in unstressed cells and the interaction was reduced in response to replication stress, (ii) RPA2 associated with the helicases through HERC2 is in its unphosphorylated form, (iii) HERC2 does not colocalize with replication stress-induced RPA2 foci, and (iv) ssDNA competes with RPA binding to HERC2 *in vitro*. Consistent with this observation, the stalled replication fork-associated proteomes did not identify HERC2 [47]. One possible scenario is that HERC2 acts on the normal replication machinery to resolve G4 structures by delivering RPA or the helicases, or both. The secondary structures of G-rich DNA occasionally impede the replication machinery, and BLM and WRN have been associated with the machinery and contribute in the removal of intramolecular G4 structures that form after dsDNA unwinding by the replicative helicase [48–50]. RPA is also capable of unfolding G4 structures in the absence of ATP and prevents G4 in the lagging strand during telomeric replication [23, 24]. HERC2 may supply RPA to the DNA locus neighboring the helicases where RPA stimulates the helicase activity by binding them directly and inhibiting the rewinding activity of the helicases, or assisting the helicases by preventing reannealing of the unfolded DNA (Figure S8B). Although a biochemical approach using purified proteins is required to elucidate the precise function of HERC2 in the process, the extremely large molecular size of HERC2 limits this at present.

It is interesting that the abundant fraction of the RPA proteins in the soluble nuclear fraction forms the complex with HERC2, and prolonged replication stress results in dissociation of RPA from the complex (Figures 2C and S3D). Given that the majority of RPA is present in the nucleosol and only one-third of RPA is associated with chromatin [51], RPA complexed with HERC2 in the soluble nuclear fraction could function as a backup system for situations with high demands for RPA, such as repair synthesis. A mechanism to save abundant cellular RPA via ATR has been reported to be critical to prevent global exhaustion of RPA that leads to a replication catastrophe in response to replication stress [52]. HERC2 may store such an RPA pool for immediate use at loci that demand BLM and WRN.

Whereas the essential role of E3 activity of HERC2 on G4 suppression *in vivo* was clear from the observation in the E3 inactivated HERC2^ΔE3/ΔE3^ cells, its molecular bases are unclear at present. As a possible event involved in the mechanism, we found that RPA accumulated in BLM or WRN complexes in the HERC2^ΔE3/ΔE3^ cells. It is possible that the E3 activity of HERC2 could be required for proper dissociation of RPA from the HERC2–helicase complexes and consequent delivery of RPA to ssDNA *in vivo*. Fine-tuning of the affinity between the helicases and RPA could be crucial in a highly repetitive DNA unfolding process, the action shared by BLM [53, 54] and other RecQ helicases [55].

It is likely that G4 accumulation itself does not harm cell viability. HERC2-depleted cells with constitutively accumulated G4 grow almost normally (Figure S6A). Furthermore, significant G4 accumulation can be detected in stomach or liver cancer tissues [56], suggesting that such cancer cells grow with increased G4. However, we show that both HERC2-depleted cells and HERC2 E3-inactivated cells were hypersensitive to the G4-stabilizers, pyridostatin and telomestatin, likely due to baseline accumulation of G4, which increases the opportunity for G4 stabilizers to bind. Telomestatin is a potent telomerase inhibitor and causes telomere shortening [57]. However, it also induces DNA damage in telomeres and other regions. Telomestatin impairs the proliferation of FANCJ-depleted cells in telomerase independent manner [6]. Importantly, G4-stabilizers are particularly toxic to BRCA-defective cells acquiring PARP inhibitor resistance [5], indicating pharmacological application exploiting G4 stabilization as a new therapeutic approach for cancers resistant to existing drugs. Although the mechanism inducing the toxicity may not be identical, HERC2 deficiency could be applicable to cancer therapy with G4 stabilizers, because HERC2 expression is frequently reduced in many types of cancers.

## MATERIALS AND METHODS

### Cell lines, Culture Conditions and Synchronization

HeLa, HCT116, U2OS and HEK293T cells were obtained from ATCC and cultured as described previously [58]. All cells were routinely monitored for mycoplasma. Cells stably expressing HERC2- or BRCA1-specific shRNA in a doxycycline-inducible manner were established by the lentiviral infection of CS-RfA-ETBsd comprising targeting sequences followed by selection with blasticidin as described [58]. The cells were treated with 1 μg/ml Dox for 48 h and subjected to individual experiments. For CRISPR/Cas9-mediated genetic engineering HCT116 cells were transfected with LentiCRISPRv2 plasmid comprising a human codon-optimized Cas9 (hSpCas9) nuclease along with a single guide RNA (sgRNA, targeting sequence: AACAGGTGTAGGACTCAGGG). Cells were selected with puromycin (1 μg/ml) and single colonies were picked up and further cultured. Mutation of HERC2 gene was confirmed by genetic sequencing and immunoblot. Cell cycle synchronization with thymidine-nocodazole block and its monitoring by flow cytometry were described previously (59).

### Chemical agents

Chemical agents used were HU (Sigma-Aldrich), MMC (Sigma-Aldrich), CPT (irinotecan hydrochloride, Sigma-Aldrich), MG132 (Calbiochem), Telomestatin (Cosmo Bio Co. LTD), Pyridostatin (Sigma-Aldrich), and BLM inhibitor (ML-216 Cayman Chemical).

### Fractionation of cellular extract

To prepare cytosolic, nuclear, and solubilized chromatin fractions, 3 × 10^7^ cells were suspended in 1 ml of the hypotonic buffer (10 mM Tris-HCl [pH 7.3], 10 mM KCl, 1.5 mM MgCl2, 10 mM β-mercaptoethanol, and 0.2 mM PMSF) and homogenized by passing through a 27-guage needle. After centrifugation at 2000 × g for 15 min at 4°C, supernatant was collected as the cytosolic fraction, and nuclei was resuspended in 700 μl of the extraction buffer (15 mM Tris-HCl [pH 7.3], 1 mM EDTA, 0.4 M NaCl, 1 mM MgCl2, 10% glycerol, 10 mM β-mercaptoethanol, and 0.2 mM PMSF) and incubated at 4°C for 30 min. The samples were then centrifuged at 20,000 × g for 30 min, and the supernatant was collected as the nuclear extract fraction. The pellet was resuspended and incubated in 1 ml of the 0.5% NP-40 buffer (50 mM Tris-HCl [pH 7.5], 0.5% Nonidet P-40, 150 mM NaCl, 50 mM NaF, 1 mM dithiothreitol, 1 mM NaVO3, 1 mM PMSF, 2 μg/ml aprotinin, 2 μg/ml leupeptin, 10 μg/ml trypsin inhibitor, and 150 μg/ml benzamidine) supplemented with 125 U/ml Benzonase nuclease (Novagen) and 2 mM Mg at 4°C for 120 min, and the reaction was stopped with 5 mM EDTA. The extract was centrifuged to isolate the chromatin-bound proteins in the soluble fraction, filtered through a 0.45-μm-pore-size filter, and used as solubilized chromatin fractions.

### Antibodies

Rabbit polyclonal antibody to C-terminal HERC2 was generated against recombinant HERC2 protein (residues 4389-4834) fused to hexa-histidine and purified by protein G agarose chromatography. The commercially available antibodies used in the study were rabbit polyclonal antibodies against HERC2 (epitope 4784-4834, Bethyl Laboratories, A301-905A), BLM (Bethyl Laboratories, A300-110A), TOP3A (Proteintech, 14525-1- AP), RMI2 (Novus Biologicals, NBP1-89962), RPA1 (Bethyl Laboratories, A300-241A), BRCA1 (Santa Cruz Biotechnology, C20), BARD1 (Bethyl Laboratories, BL518), Phospho-S33 RPA2 (Bethyl Laboratories, A300-246A), Phospho-S4/8 RPA2 (Bethyl Laboratories, A300-245A), Histone H3 (Cell Signaling, 9715), and PARP1 (Cell Signaling, 46D11) and mouse monoclonal antibodies against HERC2 (epitope 1781-1974, BD Biosciences, 17), α-tubulin (Neomarkers, DM1A, MS-581-P), RPA2 (Calbiochem, RPA34-20 NA19L), RPA3(Antibody online ABIN562685), and G4 (Absolute antibody, BG4).

### Immunoprecipitation and immunoblotting

Unless otherwise indicated cell lysates were prepared with the 0.5% NP-40 buffer followed by Immunoprecipitation and immunoblotting as described previously [60]. For the transfer of RPA from HERC2 immunocomplex to biotin-labeled DNA, cell lysates form 1 × 10^7^ HeLa cells were prepared with 1 ml of the 0.5% NP-40 buffer was clarified with centrifugation and immunoprecipitated with 2.8 μg antibody to HERC2 or control IgG covalently coupled with 0.4 mg Dynabeads (Thermo Fisher) according to the manufacturer’s instructions. The beads were washed three times with the 0.5% NP-40 buffer and once with a reaction buffer (50 mM Tris-HCl (pH 7.5), 50 mM NaCl, 5 mM MgCl2, 1 mM DTT, 5% glycerol), and resuspended in 30 μl of the reaction buffer containing indicated amount of 5’-biotin-labeled ssDNA (90nt oligo-dT, unless otherwise indicated) or dsDNA (5’- AAATCAATCTAAAGTATATATGAGTAAACTTGGTCTGACAGTTACCAATGCTTAATCAGTGAGGCACCTATCTCAGCGATCTGTCTATTT-3’, the sequence complementary to positions 1932-2022 of the pBluescript replicative form I DNA used in a previous study [61]), incubated for 1 min at 25 °C/950 r.p.m on ThermoMixer (Eppendorf). The reaction was terminated by moving the reaction tubes on a magnetic rack. The elution was incubated with 10 μl of 50% volume of prewashed StrepTactin beads (IBA), incubated for 10 min at room temperature, washed with the 0.5% NP-40 buffer. The StrepTactin beads were then boiled with NuPAGE sample buffer (Thermo Fisher). The remained dynabeads were washed with the 0.5% NP-40 buffer, eluted with 30 μl of elution buffer (0.5 M NH_4_OH, 0.5 mM EDTA; Thermo Fisher) and boiled with NuPAGE sample buffer according to the manufacturer’s instructions. Boiled samples were subjected to immunoblotting.

### Affinity purification of HERC2 complex and Mass Spectrometry

Cell lysates form 3 × 10^7^ HeLa cells and HCT116 cells treated or not with MMC (0.5 μg/ml) for 16 h or MG132 (5 μM) for 12 h were prepared with 3 ml of the 0.5% NP-40 buffer. After clarify with centrifugation the lysates were immunoprecipitated with antibody to HERC2 or control IgG covalently coupled with Dynabeads (Thermo Fisher), washed, eluted with an elution buffer (0.5 M NH4OH, 0.5 mM EDTA) according to the manufacturer’s instructions. Samples were concentrated with an evaporator and subjected to 4–12% NuPAGE Bis-Tris gels (Life Technologies) with a short run (1 cm). The gels were stained with Bio-Safe Coomassie Stain (Bio-Rad). After the gels were extensively washed with Milli-Q water (Millipore), the gel regions of entire lane were excised, cut into 1 mm^3^ pieces, destained for 1 h with 1 ml 50 mM ammonium bicarbonate (AMBC)/30% acetonitrile (ACN) with agitation and then further washed for 1 h with 1 ml 50 mM AMBC/50% ACN. Finally, a 100% ACN wash was performed to ensure complete gel dehydration. Trypsin solution (Promega, 20 ng/μl in 50 mM AMBC/5% ACN) was subsequently added to the gel pieces at approximately equivalent volumes and incubated on ice for 30 min. Another small volume of trypsin solution was added to the gel samples and incubated at 37°C overnight. Digests were extracted by addition of 100 μl 50% ACN/0.1% trifluoroacetic acid (TFA) for 1 h with shaking. The peptides were recovered into fresh Eppendorf tubes, and an additional extraction step was performed with 70% ACN/0.1% TFA for 30 min. The extracted peptides were concentrated using a speed-vac, and subjected to shotgun MS analysis on a Q Exactive mass spectrometer coupled with an EASY-nLC 1000 liquid chromatograph and nanoelectrospray ion source (Thermo Scientific). The mobile phases were 0.1% formic acid (FA) in water (Solvent A) and 0.1% FA in 100% ACN (Solvent B). Peptides were directly loaded onto a C18 analytical column (ReproSil-Pur 3 μm, 75 μm inner diameter and 12 cm length, Nikkyo Technos) and separated using a 80 min two-step gradient (0–35% Solvent B for 70 min and 35–100% for 10 min) at a constant flow rate of 300 nl/min. The Q Exactive was operated in the data-dependent MS/MS mode, using Xcalibur software, with survey scans acquired at a resolution of 70,000 at m/z 200. The top 10 most abundant isotope patterns with charge 2–5 were selected from the survey scans with an isolation window of 2.0 m/z, and fragmented by HCD with normalised collision energies of 28. The maximum ion injection times were 60 ms for both survey and MS/MS scans, and the AGC values were set to 3 × 10^6^ and 5 × 10^5^ for the survey and MS/MS scans, respectively. Ions selected for MS/MS were dynamically excluded for 10 sec. Proteome Discoverer software (ver. 1.3, Thermo Scientific) was used to generate peak lists. The MS/MS spectra were searched against a Swiss-Prot database (version 2012_10 of UniProtKB/Swiss-Prot protein database) using the MASCOT search program (Matrix Science). The precursor and fragment mass tolerances were set to 10 ppm and 20 mmu, respectively. Maximum missed cleavage site of trypsin was set to two. Acetylation of N-terminal protein, oxidation of methionine, ubiquitylation/acetylation of lysine, phosphorylation of serine/threonine, pyroglutamylation of N-terminal glutamine were set as variable modifications for database searching. Peptide identification was filtered at a 1% false discovery rate.

### siRNAs, Plasmids, and Transfections

siRNA oligonucleotides targeting BLM (HSS101023), WRN (S14908), TOP3A (HSS110902), RPA1 (S12129), and RPA2 (S12131), and the non-targeting control siRNA (AM4635) were purchased from Ambion. RNA duplexes (final concentration, 10 nM) were transfected into the cells with Lipofectamine RNAiMAX (Invitrogen) and analyzed 48 h after the transfection. Vectors for the doxycycline-inducible expression of shRNAs were generated with insertion of oligos containing the sequence specific to HERC2 (5’- GAAGGTGGCTGTTCACTCA-3’) into the lentiviral vector plasmid CS-RfA-ETBsd via Gateway LR recombination from pENTR4-H1tetOx1. CS-RfA-ETBsd and pENTR4-H1tetOx1 are gifts from Dr. Hiroyuki Miyoshi and Dr. Atsushi Miyawaki, RIKEN BioResource Center.

### Gel Filtration Analysis (Size Exclusion Chromatography)

Nuclear extracts (2 mg/0.5 ml) were applied to Superose 6 10/300 GL column (GE Healthcare) and fractionated using an AKTA Purifier (GE Healthcare) liquid chromatography system with a flow rate of 0.4 ml/min in buffer consisting of 50 mM Tris-HCl (pH 7.4), 1 mM EDTA, 0.1% CHAPS, 1 mM DTT, and 150 mM NaCl. 500 μl fractions were collected after 5 ml and analyzed by western blot analysis. The approximate molecular weight was estimated with Gel Filtration Calibration kit (GE Healthcare).

### Immunofluorescence microscopy

HERC2/RPA2 staining was performed using cold methanol and acetone as described previously [35]. For staining of G4, mixed methanol:acetic acid (3:1) solution was used as described elsewhere [45] followed by 0.5% Triton X-100 in PBS with 200 μg/ml RNase A. Subsequently, the cells were washed, blocked with 3% goat serum and 0.1% Triton X-100, and labeled with primary and fluorescence-labeled secondary antibodies. The slides were mounted with ProLong Gold Antifade Mountant with DAPI (Invitrogen) and examined with a confocal laser-scanning microscope (LSM 510, Carl Zeiss, Germany). G4 nuclear foci were mechanically counted using the Cellomics Image Analyzer (Thermo Fisher).

### Clonogenic survival assay

Cells were seeded at a concentration of 500 cells/well in 6-well plates and after 6 h different concentrations of agents were added. After 24 h of incubation, the cells were washed and further cultured in fresh medium without the chemicals for 14 days. The cells were then fixed and stained with crystal violet. The colonies were scanned and counted using an ImageQuant LAS-4000 instrument (GE Healthcare).

### TCGA data analysis

Normalized gene expression data (dataset: gene expression RNAseq - UNC RSEM norm_count) of HERC2 and clinical information in TCGA Pan-Cancer data were downloaded using the UCSC Xena browser (http://xena.ucsc.edu/). Values are log2(x+1) transformed RSEM values. p-values for differential expression between cancer (reffered as ‘Primary Tumor’ in the dataset) and normal (reffered as ‘Solid Tissue Normal’) were calculated by two-tailed Student’s t test for each type of cancer. The package of ggplot2 for R were used to generate a box plot.

## AUTHOR CONTRIBUTIONS

W.W, assisted by Y.T, performed most of the experimental works. N.R characterized HERC2 antibody and assisted in CRISPR/Cas9 experiments. J.T and N.A, assisted by Y. S, performed the mass-spectrometric analysis. Y.L contributed to immunofluorescence and clonogenic survival assays. R.M analyzed TCGA data. H.N performed gel filtration analysis. Y.M and N.S supervised the project. Y.S and K.T supervised the mass-spectrometric analysis and provided advice. T.O designed and supervised experiments and wrote the manuscript. All authors critically read the paper.

## ACKNOWLEDGMENTS

We thank T. Fukuda for advice in clonogenic survival assay and S. Hanaki for technical support. We are grateful to B. Shiotani and M. Nakanishi for helpful discussion. This work was supported by MEXT/JSPS KAKENHI (grant numbers 17H03585, JP26290042 and JP 24112005 to T.O, JP26460402 to W.W., JP24112008 to Y.S., and JP26000014 to K.T.) and the Japan Agency for Medical Research and Development (grant number 16ck0106085h0003) to T.O.

